# Patterns of mesozooplankton community composition and vertical fluxes in the global ocean

**DOI:** 10.1101/2021.07.20.452978

**Authors:** Yawouvi Dodji Soviadan, Fabio Benedetti, Manoela C. Brandão, Sakina-Dorothée Ayata, Jean-Olivier Irisson, Jean Louis Jamet, Rainer Kiko, Fabien Lombard, Kissao Gnandi, Lars Stemmann

## Abstract

Vertical variations in physical and chemical conditions drive changes in marine zooplankton community diversity and composition. In turn, zooplankton communities play a critical role in regulating the transfer of organic matter produced in the surface ocean to deeper layers. Yet, the links between zooplankton community composition and the strength of vertical fluxes of particles remain elusive, especially on a global scale. Here, we provide a comprehensive analysis of variations in zooplankton community composition and vertical particle flux in the upper kilometer of the global ocean. Zooplankton samples were collected across five depth layers and vertical particle fluxes were assessed using continuous profiles of the Underwater Vision Profiler (UVP5) at 57 stations covering seven ocean basins. Zooplankton samples were analysed using a Zooscan and individual organisms were classified into 19 groups for the quantitative analyses. Zooplankton abundance, biomass and vertical particle flux decreased from the surface to 1000m depth at all latitudes. The zooplankton abundance decrease rate was stronger at sites characterised by oxygen minima (< 5µmol O_2_.kg^−1^) where most zooplankton groups showed a marked decline in abundance, except the jellyfishes, molluscs, annelids, large protists and a few copepod families. The attenuation rate of vertical particle fluxes was weaker at such oxygen-depleted sites. Canonical redundancy analyses showed that the epipelagic zooplankton community composition depended on the community structure of surface phytoplankton and the quantity and the quality of the produced particulate organic matter. We provide a consistent baseline of plankton community structure together with estimates of vertical flux and a contribution to our understanding of global zooplankton dynamics in the upper kilometer of the global ocean. Our results further suggest that future changes in surface phytoplankton composition and mesopelagic oxygen loss might lead to profound changes in zooplankton abundance and community structure in both the euphotic and mesopelagic ocean. These changes may affect the vertical export and hereby the strength of the biological carbon pump

## Introduction

The upper kilometer of the ocean constitutes a habitat where most of the organic carbon produced by the phytoplankton in the epipelagic layer (0-200m) sinks into the mesopelagic layer (200-1000m) while being progressively consumed and respired. Within the wide size spectrum of planktonic organisms (0.02µm-2m), mesozooplankton (0.2-20mm) are a pivotal component of marine trophic webs impacting the Biological Carbon Pump (BCP) through their feeding, vertical migration, and the production of faecal pellets (Steinberg and Landry, 2017). In a context of global climate change, zooplankton communities experience increasingly stressful conditions through global warming, ocean acidification and deoxygenation (Oschlies et al., 2018; Schmidtko et al., 2017), enhanced water column stratification in the open ocean and modifications in phytoplankton production and community structure (Coma et al., 2009; Kwiatkowski et al., 2019; Richardson, 2008). Long term field surveys have shown how shifts in climatic conditions lead to large shifts in surface zooplankton composition (Beaugrand et al., 2019). Future climate warming could further reduce macronutrient supplies in the upper ocean and therefore decrease phytoplankton biomass in the tropical open ocean or can cause positive trophic amplification in the polar ocean (Chust et al., 2014). Such changes in biomass may be amplified in the zooplankton through trophodynamic effects, which could greatly alter the fluxes of organic matter into the deeper mesopelagic layers. Within the latter, modifications of the quantity and quality of the vertical particle flux produced in the surface layers could trigger substantial changes in mesozooplankton abundance and composition. In addition, the expansion of Oxygen Minimum Zones (OMZ) (Schmidtko et al., 2017; Stramma et al., 2010) may further constrain the spatial distribution of the numerous zooplankton taxa that are sensitive to dissolved oxygen levels (Ekau et al., 2010; Kiko et al., 2020; Kiko and Hauss, 2019; Seibel, 2011; Wishner et al., 2020, 2018). Therefore, understanding the variations of zooplankton biomass and diversity in the epi and mesopelagic is essential to better understand the impact of global climate change on the properties of marine ecosystems.

However, mesopelagic zooplankton communities remain critically under-sampled compared to the epipelagic ones, as most collections is in the first 200m of the water column (Everett et al., 2017). The mesopelagic is still considered a “dark hole” (Robinson et al., 2010; St John et al., 2016) as the gaps in sampling lead to critical gaps in knowledge regarding the functioning of mesopelagic ecosystems and how they are controlled by the organic matter fluxing from the euphotic zone. Local to regional field studies reported that the abundance of mesozooplankton decreases exponentially with depth, in parallel with substantial changes in species diversity and genus composition that remain poorly resolved (Bode et al., 2018; Brugnano et al., 2012; Hernández-León et al., 2020; Koppelmann et al., 2005; Kosobokova and Hopcroft, 2010; Yamaguchi et al., 2004). Some studies reported increases in diversity with depth, with maxima in the meso- to bathypelagic layers (Bode et al., 2018; Gaard et al., 2008; Kosobokova and Hopcroft, 2010; Yamaguchi et al., 2004), whereas other studies reported either the opposite diversity pattern or even an absence of vertical patterns due to large spatio-temporal variability (Gaard et al., 2008; Hidalgo et al., 2012; Palomares-García et al., 2013; Wishner et al., 2008). Such mismatch between observations might be related to the wide range of environmental conditions sampled using a variety of sampling gears that make inter-comparisons difficult. One recent global analysis of zooplankton vertical distribution in inter-tropical regions showed that in this narrow latitudinal band the vertical dimension was the main structuring pattern suggesting that midwater processes mediated by zooplankton may not change between oceans at those latitudes (Fernández de Puelles et al., 2019). How these changes relate to the vertical flux and also across the full latitudinal band is poorly understood because plankton observations and biogeochemical studies have usually been carried out by distinct scientific communities. Recent interdisciplinary cruises have taken place, providing more integrated knowledge on the importance of zooplankton to carbon sequestration (Guidi et al., 2016; Kiko et al., 2020; Steinberg and Landry, 2017; Stukel et al., 2019). No global study has yet been able to analyse community-level variations in zooplankton abundances, biomass and community composition in conjunction with assessments of vertical particle flux in the upper kilometre of the global ocean.

Here, a homogeneous dataset of community-level zooplankton images generated from the TARA Oceans expeditions is explored together with associated measurements of their contextual environmental conditions to: (i) test whether zooplankton communities present consistent vertical variations in abundance and composition across the oceans and in different latitudinal bands, and (ii) investigate the main drivers of such vertical gradients and how they are coupled to the surface phytoplankton and estimates of particle flux. The TARA Oceans expedition collected over 35000 samples encompassing the whole plankton community (i.e., from viruses to jellyfish) across the global ocean from 2009 to 2013 (Karsenti et al., 2011). Previous studies that analysed the TARA Oceans imaging dataset explored the global latitudinal gradients in richness and composition for the whole plankton community focusing on surface waters (Ibarbalz et al., 2019; Brandao et al., in press). Here, we aim to describe the community composition of mesozooplankton down to 1000m depth on a global scale. As the imaging techniques used here cannot achieve a species-level identification of the zooplankton community composition, our approach is rather oriented towards larger taxonomic and /or functional groups that may be associated with broad ecological and biogeochemical functions.

## Materials and methods

### Sampling sites and environmental variables

TARA Oceans took place between 2009 and 2013. Among the 210 stations that were sampled, 57 stations covering 7 ocean basins (Fig. 1) were sampled with a Multinet (Pesant et al., 2015; Roullier et al., 2014), a sampling device with five nets that allows for depth-stratified sampling (see below).

**Fig. 1:**
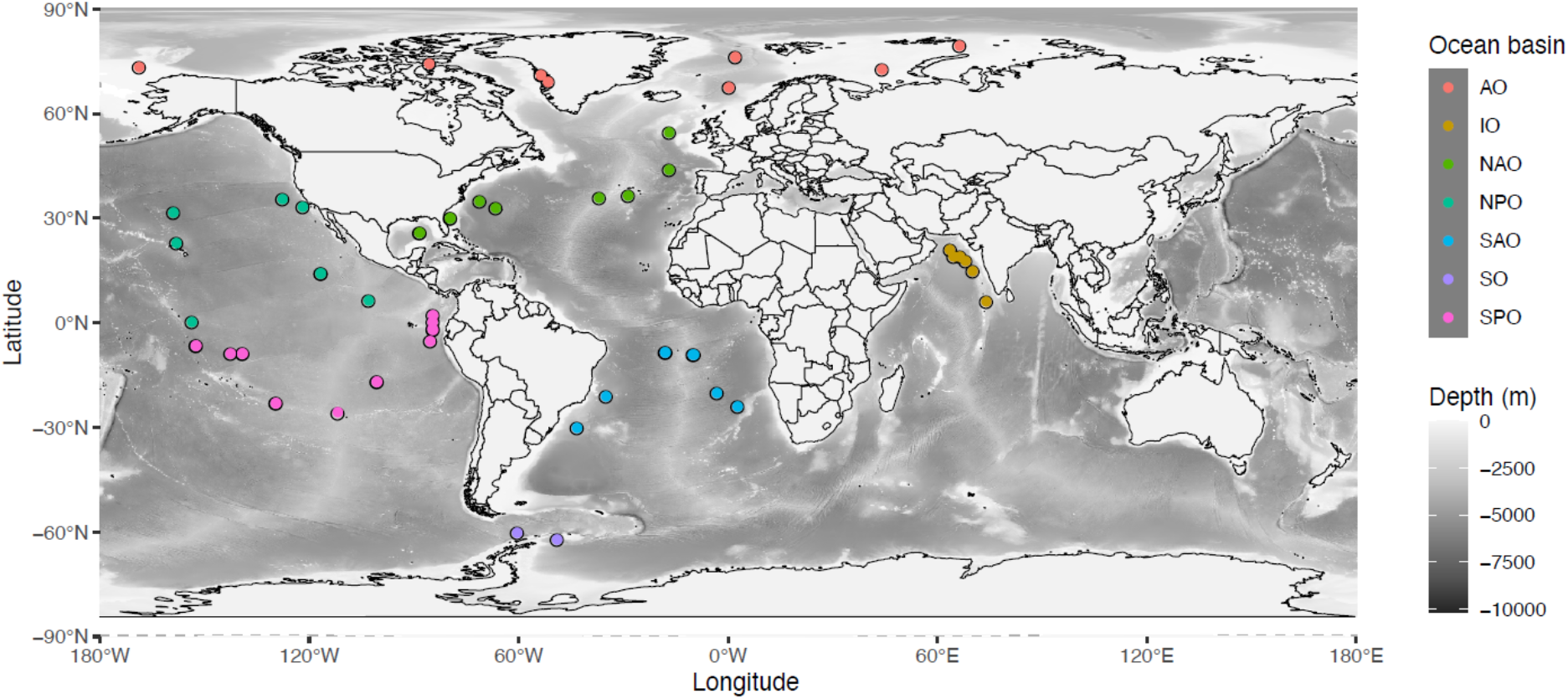
Location of the 57 stations sampled with the Multinet, grouped by ocean basin: AO=Arctic Ocean, IO=Indian Ocean, NAO=North Atlantic Ocean, NPO=North Pacific Ocean, SAO=South Atlantic Ocean, SO=Southern Ocean or Austral Ocean, SPO=South Pacific Ocean.

A CTD rosette equipped with optical sensors was deployed to measure the physico-chemical parameters within the water column. Temperature and conductivity were measured from the surface to a maximum of 1300 m depth using a Seabird 911 CTD mounted on a Sea-Bird Carousel sampler with 10 Niskin bottles. The following additional sensors were mounted to measure optical properties related to relevant biogeochemical variables: fluorometer (Wetlab ECO-AFL/FL model), dissolved oxygen sensor (model SBE 43), nitrate sensor (ISUS with a maximum rating depth of 1000m Satlantic SA), a 25 cm transmissometer for particles 0.5–20 µm (WETLabs), a one-wavelength backscatter meter for particles 0.5–10 µm (WETLabs), and an Underwater Vision Profiler 5 (UVP5) for particles >150 µm and zooplankton >600 µm (Hydroptic). Assuming that particle sinking speed increases with size, those particles detected through the backscattering will be referred to as suspended particulate matter (SPM, particles < 10µm in Equivalent Spherical Diameter), while the ones detected by the UVP5 (>150µm in Equivalent Spherical Diameter) will be referred as particles. Vertical particle mass flux (in mg Dry Weight m^-2^d^-1^)) was calculated from the particle size spectra detected by the UVP5 as in (Guidi et al., 2008). Based on the High Pressure Liquid Chromatography (HPLC) analysis of water collected with Niskin bottles, we used the method of Uitz et al. (2006) to estimate the contribution (%) of three pigment size classes (microphytoplankton, nanophytoplankton, and picophytoplankton; *f__micro_*, *f__nano_*, and *f__pico_*, respectively) to total phytoplankton biomass in the epipelagic layer. The median value of all hydrological and optical data together with imaging data were calculated for each of the five horizontal layers sampled by the Multinet for future data processing. The samples were classified as anoxic, hypoxic and normoxic according to the oxygen minimum value measured within the towed layer. We used a threshold of 5 µ*mol kg*^-1^ to classify the stations as anoxic (Roullier et al., 2014) and 58.5 µ*mol kg*^-1^(Bode et al., 2018) to classify them as hypoxic.

### Zooplankton sampling, digitization, biomass estimates

A Hydrobios Multinet (with a 300µm mesh and an aperture of 0.25m^2^) was used to sample zooplankton (Roullier et al., 2014; Pesant et al., 2015) in five distinct water layers ranging from the surface to occasionally 1300 m depth. The five depth layers were locally defined as a function of the vertical structure of the water column according to the profiles of temperature, salinity, fluorescence, nutrients, oxygen, and particulate matter (Pesant et al., 2015). The Multinet was equipped with a flowmeter to measure the volume of seawater filtered by each net tow (Pesant et al., 2015). Day and night net tows were conducted at ten stations. Sampling at the other stations occurred at day or night, depending on cruise schedule and operational constraints. Once collected, the samples were preserved in a solution of buffered formaldehyde-Borax solution (4%). In the laboratory, the samples were rinsed and split with a Motoda box (MOTODA, 1959). The final split was analysed with the Zooscan imaging system (Gorsky et al., 2010) which allowed a rapid and semi-automatic analysis of zooplankton samples. In total, the samples comprised nearly 400,000 images of living zooplankton and detritus. These images were imported into Ecotaxa, an online platform which allows an automatic prediction of the taxonomic classification of each single image followed by a manual validation/correction. The organisms were then identified manually down to the order, sometimes to the family and rarely down to the genus level. All copepods were sorted at the family level apart from the smallest copepods that cannot be recognised at the family level from the image. They were all grouped into one category called Other-copepoda or other-Calanoida if their morphological features allowed classifying them as Calanoida. This initial sorting allowed classifying zooplankton into 119 taxa. As many taxa showed a very small contribution to total zooplankton abundance, the 119 taxa were grouped into 19 taxonomic groups (Table 1). Those include all the major zooplankton groups that are frequently observed in the oceans. To investigate vertical patterns in mesozooplankton abundance, these 19 groups were further aggregated into eight groups representing a combination of taxonomic and functional classification (Table 1).

**Table 1:** List of the 19 taxa kept for the RDA analysis.

| Taxonomic groups | Sub groups |
| --- | --- |
| Protista | Protista includes unrecognized protista on the images |
| Annelida | Images of all Annelida |
| Chaetognatha | Images of all Chaetognatha |
| Tunicata | Images of all Gelatinous filter feeders |
| Cnidaria | Images of all gelatinous carnivores belonging to Cnidaria |
| Mollusca | Images of all Pteropoda |
| Cladocera | Images of all Small Cladocera |
| Eumalacostraca | Images of all Decapoda but those distinguish at detailed taxonomic level |
| Euphausiacea | Images of all Euphausiacea |
| Amphipoda | Images of all Amphipoda |
| Ostracoda | Images of all Ostracoda |
| Crustacean_larvae | Images of all crustacean larvae |
| Other_Copepoda | Images of all Copepoda but those distinguish at detailed taxonomic level |
| Calanoida | Images of all Calanoida but those distinguish at detailed taxonomic level |
| Oithonidae | Images of all Oithonidae |
| Corycaeidae | Images of all Corycaeidae |
| Oncaeidae | Images of all Oncaeidae |
| Metridinidae | Images of all Metridinidae |
| Calanidae | Images of all Calanidae |

Once the zooplankton images were sorted, Ecotaxa enabled us to extract the concentration and the biovolume of each mesozooplankton group at every station and for every net tow, while accounting for the Motoda fraction and the volume sampled. The biovolume was computed for each individual zooplankton using the minor and major length axes assuming a prolate ellipsoidal shape (Gorsky et al., 2010). The biomass was calculated for the 8 large groups using the equations for the different taxa :

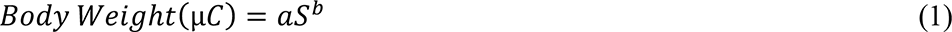

where *S* is body area in *mm*^2^. Taxon-specific area-to-dry mass conversion factors (Lehette and Hernandez-Leon, 2009) and dry mass to carbon (C) conversion factors (Kiørboe, 2013) were used to calculate the biomass and C content of each zooplankton organism scanned. Taxonomic units and biomass conversion factors used are listed in Table 2. For large protists the conversion factor was adjusted to 0.08 *mgC mm*^-3^ (Biard et al., 2016).

**Table 2:** Coefficient factors used for equation 1 and that were taken from observed allometric relationships between body area and individual dry mass (Lehette et al., 2009). The conversion factors to carbon are taken from Kiorboe 2013.

| Zooplankton category | Biomass Exponent (b) | Biomass Multiplier (a) | DW to C |
| --- | --- | --- | --- |
| Copepods | $1.59 \pm 0.03$ | 45.25 | 0.48 |
| Chaetognaths | $1.19 \pm 0.13$ | 23.45 | 0.367 |
| Decapods | $1.48 \pm 0.05$ | 49.58 | 0.419 |
| Cnidaria | $1.02 \pm 0.38$ | 43.17 | 0.132 |
| Tunicata | $1.24 \pm 0.08$ | 4.03 | 0.103 |
| Pteropods | $1.54 \pm 0.03$ | 43.38 | 0.289 |
| Protists* |  | 0.08 mgC mm<sup>-3</sup> |  |
| Other<br>(Annelids, Cladocera,<br>Ostracoda) | $1.54 \pm 0.03$ | 43.38 | 0.289 |

\* for the protist we used a conversion factor between biovolume and biomass (Biard et al., 2016)

Shannon diversity index (*H*^’^) was calculated based on the relative abundances of the 119 taxa for each sample as follows:

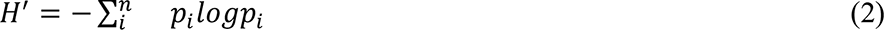

where *p_i_* is relative abundance of each taxa in one sample and log is the natural logarithm.

### Analyzing zooplankton and particles vertical distributions

Vertical attenuation rates of zooplankton (abundance and biomass) and estimated particle fluxes were estimated, from the five sampled layers for zooplankton and from the 5 meter resolution profile of estimated vertical flux using a linear regression of the log-log (i.e. natural logarithm) with the following equation :

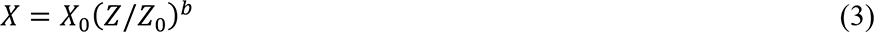

where *X* represents the zooplankton abundance, the zooplankton biomass or the particle vertical flux at the depth level *Z*, *X*_O_the zooplankton biomass or abundance and vertical particle flux at the depth *Z*_O_ (chosen as median depth of the surface net) and *b* the slope taken as a proxy of the attenuation rate of zooplankton biomass zooplankton abundance or particle flux. In the rest of the manuscript, A_zoo represents the slope *b* of vertical profile for zooplankton abundance, B_zoo the slope *b* of vertical profile for zooplankton biomass, A_flux the slope *b* of vertical profile for particle flux, and P_flux1, P_flux2 and P_flux3 the particle flux in respectively the epipelagic, upper and lower mesopelagic. To analyse latitudinal patterns in attenuation rates, the slope values were separated into three latitudinal bands based on the latitudinal position of their corresponding sampling stations: intertropical (0°-30°), temperate (30-60°) and polar (60°-90°). The intertropical stations gathered both OMZ and non-OMZ stations, which allowed us to analyse the effect of oxygen depletion on zooplankton and particles. Non-parametric variance analyses (Kruskal and Wallis tests) were performed to test for differences in slope values (i.e. zooplankton and particles attenuation rates) between latitudinal bands.

### Multivariate analysis of community composition

To explore the response of zooplankton community composition to environmental drivers across depth layers, the non-interpolated abundances of the 19 taxonomic groups mentioned above were aggregated into three layers: the epipelagic layer (0-200m), the upper mesopelagic layer (200-500m) and the lower mesopelagic layer (500-1000m). To analyse separately the three depth layers, the samples collected in overlapping layers (18.59% of the total samples) were not included in the statistical analysis (Table S1). To characterise the environmental conditions of each layer at each sampling station the median values of the following contextual environmental variables were used: temperature (T), salinity (S), oxygen (O_2_), nitrate concentration (NO_3_), chlorophyll *a* concentration (Chl_*a*), phytoplankton size fractions (*f__micro_*, *f__nano_*, and *f__pico_*), concentration of suspended particles (SPM) and particle flux (P_Flux). The measurements of all these environmental variables are available on PANGAEA (https://doi.org/10.1594/PANGAEA.840721).

To estimate the strength of the Diel Vertical Migration (DVM) at 10 stations, pairwise Wilcoxon tests were performed to compare in each layers the abundance and biomass of each taxa between day and night. For those 10 same pairs of stations, we used an analysis of similarities (ANOSIM) to test for significant variations in community composition between day and night samples. The ANOSIM tested whether the inter-groups difference (day and night groups) was higher than the intra-groups difference, by providing an R coefficient. An R coefficient close to one suggests dissimilarity between groups, while R value close to zero suggests an even distribution of high and low ranks within and between groups. An R value below zero suggests that dissimilarities are greater within groups than between groups (Clarke and Gorley, 2001). ANOSIM tests were performed within each layer using log-transformed (where log is the natural logarithm) abundances and Bray-Curtis distance among stations.

For each depth layer, a canonical redundancy analysis (RDA) was performed based on the abundances of the 19 mesozooplankton groups and the above-mentioned environmental variables to explore the explanatory power of these variables in structuring the mesozooplankton community. The RDA is an extension of the multiple regression analysis applied to multivariate data (Legendre and Legendre, 1998). It allows representing the response variables (abundances of the 19 mesozooplankton groups) in a “constrained” reduced space, i.e., constructed from the explanatory variables (the environmental variables). For each RDA, the following variables were used as “supplementary variables” of the analysis in order to visualize their correlation with the environmental structuring of the mesozooplankton community (i.e., to visualise their position in the RDA space): attenuation of particle flux (A_flux), attenuation of zooplankton abundance (A_zoo), attenuation of zooplankton biomass (B_zoo) and the Shannon index (*H’*). Beforehand, a Hellinger transformation was performed on the mesozooplankton abundances. A preliminary RDA based on all samples together showed a very strong effect of depth on mesozooplankton community composition (Fig. S1). Therefore, to avoid such a strong effect of depth on the community composition analysis, three separate RDAs were performed on each of the three layers defined above. Significant axes were identified using the Kaiser-Guttman criterion (Legendre and Legendre, 1998).

Data manipulation and statistical analyses were performed with Matlab 2018b (MATLAB 9.5) for the vertical profiles of abundance and biomass and statistical test (Wilcoxon test, Kruskal-wallis test), R environment v3.5.1 (using the following packages: vegan version 2.5-5, ggplot2 version 3.1.1, ggrepel version 0.8.0 and ggfortify version 0.4.7) for the redundancy analysis and PRIMER6 (Version 6.1.12) and PERMANOVA+ (Version 1.0.2) for the ANOSIM test.

## Results

### Day night variability

In each layer, the comparison between day/night of the taxonomic groups indicated that only a few groups (Euphausiidae, Metridinidae, Corycaeidae and Cnidaria for abundance; Eumalacostraca and Ostracoda for biomass) showed significant differences in either the surface layer or the upper mesopelagic layer (Table S1). Nonetheless, the ANOSIM found no significant differences in the community composition between the day and night samples at the community-level (R0-200m = −0.039; R200-500m = −0.008; R500-1100m = −0.006). The same analyses based on biomass also indicated no significant differences (R0-200m = 0.018; R200-500m = −0.051; R500-1100m = 0.087). As a consequence, all further analyses were carried out without making any distinctions between day and night samples.

### Vertical patterns in zooplankton total abundance and biomass and particle flux

On a global scale, zooplankton abundance and biomass decreased exponentially with depth (Fig. 2) in the different latitudinal bins. Abundance and biomass decrease rate with depth and were correlated (r2=0.342, p=4.6 10^-5^) but the biomass attenuation rate estimates were systematically lower than the attenuation based on the abundance profiles. On average, polar waters showed increased zooplankton abundance and biomass compared to the stations located in the tropics. In the epipelagic layer, abundances and biomass ranged from 1 to 5000 ind m^-3^ and 0.05 to 200 mg C m^-3^ while in the mesopelagic they were reduced to 0.05 to 450 ind m^-3^ and 0.005 to 40 mg C m^-3^. Copepods were the most abundant being 85% and 65% of the abundance and biomass in the epipelagic, 85 and 76% in the upper mesopelagic and 95% and 97% in the lower pelagic (Table 3). The estimated vertical flux also decreased with depth in all latitudinal bands. On average polar waters showed higher fluxes compared to the stations located in the tropics.

**Fig. 2:**
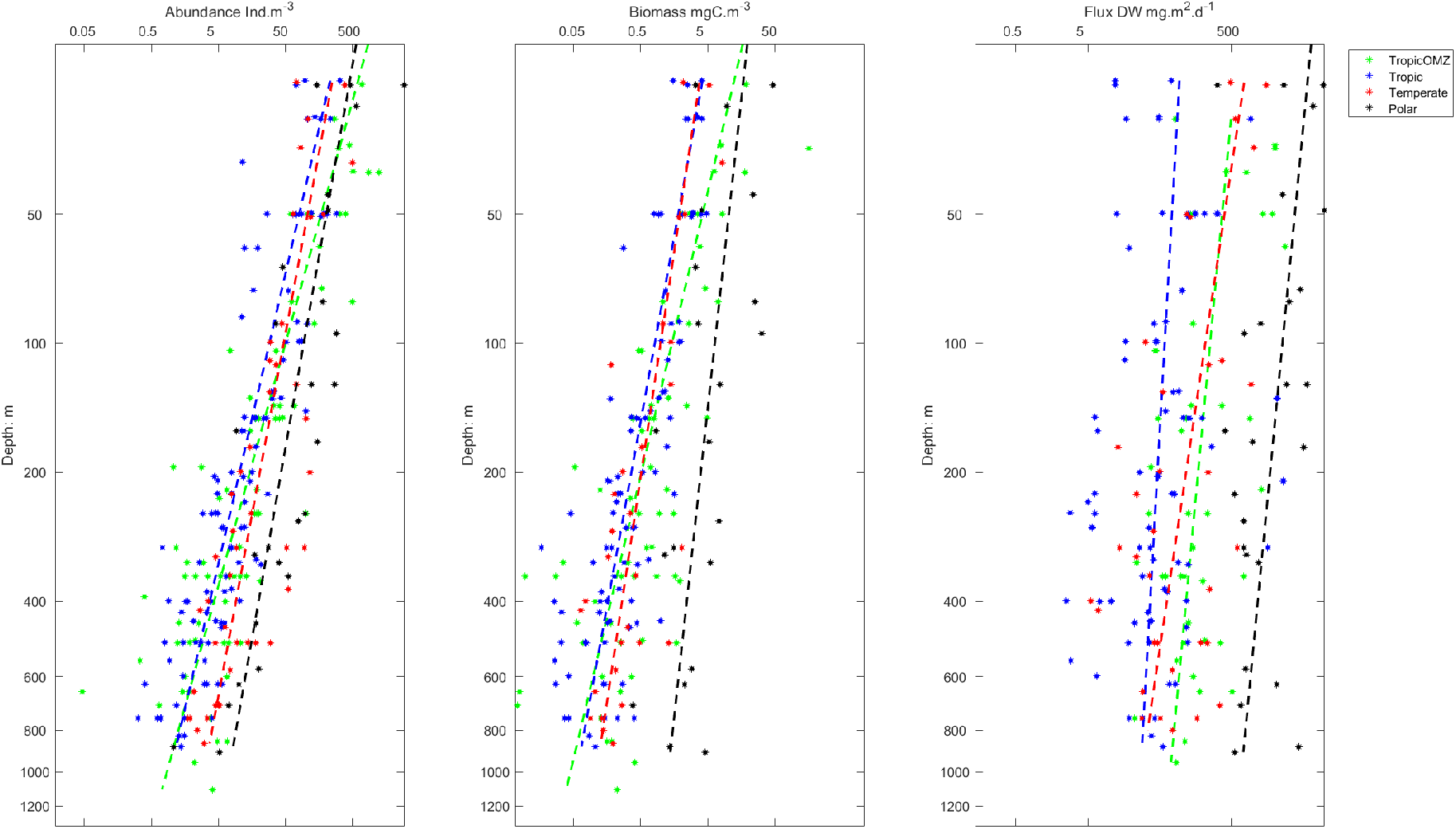
Vertical distribution of zooplankton biomass, abundance and particle flux in different latitudinal bands. Dotted lines represent the fitted linear regression for each latitudinal band (equation 3). The linear fits to the data point are given in table 4.

**Table 3:** Summary of the main mesozooplankton groups global proportion of abundance and biomass:

| Taxa<br>Proportion(%) | Epipelagic |  | Upper Mesopelagic |  | Lower Mesopelagic |  |
| --- | --- | --- | --- | --- | --- | --- |
|  | Abundance | Biomass | Abundance | Biomass | Abundance | Biomass |
| All copepoda | 84.89 | 65 | 84.80 | 76.39 | 94.5 | 96.5 |
| Chaetognatha | 3.67 | 8.39 | 2.44 | 3.48 | 2.06 | 2.53 |
| Crust decapoda | 2.9 | 18.9 | 0.36 | 14.98 | 0.00 | 0.00 |
| Cnidaria | 0.7 | 0.96 | 0.19 | 0.24 | 0.00 | 0.00 |
| Tunicata | 0.5 | 0.02 | 0.08 | 0.001 | 0.1 | 0.00 |
| Protista | 1.5 | 0.5 | 0.97 | 0.19 | 0.9 | 0.09 |
| Pteropoda | 1.2 | 0.6 | 0.41 | 0.03 | 0.00 | 0.00 |
| Other<br>(annelids, cladocera,<br>ostracoda) | 4.2 | 3.09 | 10.05 | 3.23 | 2.38 | 0.87 |

**Table 4:** Summary of the vertical attenuation rate (b: median and IC: confidence Interval) of mesozooplankton abundance and biomass and particle vertical flux (b: median value and IC: confidence Interval).

|  | Zooplankton<br>abundance<br>(Ind. m<sup>-3</sup>) |  | Zooplankton biomass<br>(mgC. m<sup>-3</sup>) |  | Particle vertical flux |  |
| --- | --- | --- | --- | --- | --- | --- |
|  | b | IC (95%) | b | IC (95%) | b | IC (95%) |
| ANOXIA_TROPIC | -1.82 | -2.13 - -1.13 | -1.3 | -2.9 - -0.75 | -0.26 | -0.77- -0.19 |
| TROPIC | -1.39 | -2.46 - -0.69 | -1.04 | -1.92 - -0.58 | -0.42 | -0.78- -0.21 |
| TEMPERATE | -1.19 | -1.56 - -0.64 | -0.89 | -1.26 - -0.57 | -0.50 | -0.84- -0.11 |
| POLAR | -1.18 | -1.71 - -0.96 | -0.83 | -1.41 - -0.74 | -0.78 | -4.51- -0.36 |

The attenuation rates of zooplankton abundance and biomass with depth were stronger than the attenuation rates of the vertical particle flux (Fig. 3A, B and C, Table 4). The decrease rates in zooplankton vertical abundance and biomass are more pronounced in OMZ stations compared to the non-OMZ stations. Yet, such difference was found to be significant only when the vertical decrease rates were calculated from abundances. In general, zooplankton attenuation rates decreased with latitude whereas the attenuation rates of particle fluxes increased with latitude (Fig. 3, Table 3). A non-parametric variance analysis (Kruskal-Wallis test) of the attenuation rate revealed significant differences in the attenuation rates of zooplankton abundances between the anoxic tropical stations and the temperate and polar ones, but not with the other tropical stations (Table 4). No statistical difference between regions was found. The attenuation rate of vertical flux was weaker at the OMZ sites compared to the non OMZ ones but this difference was not significant (Fig. 3B).

**Fig. 3:**
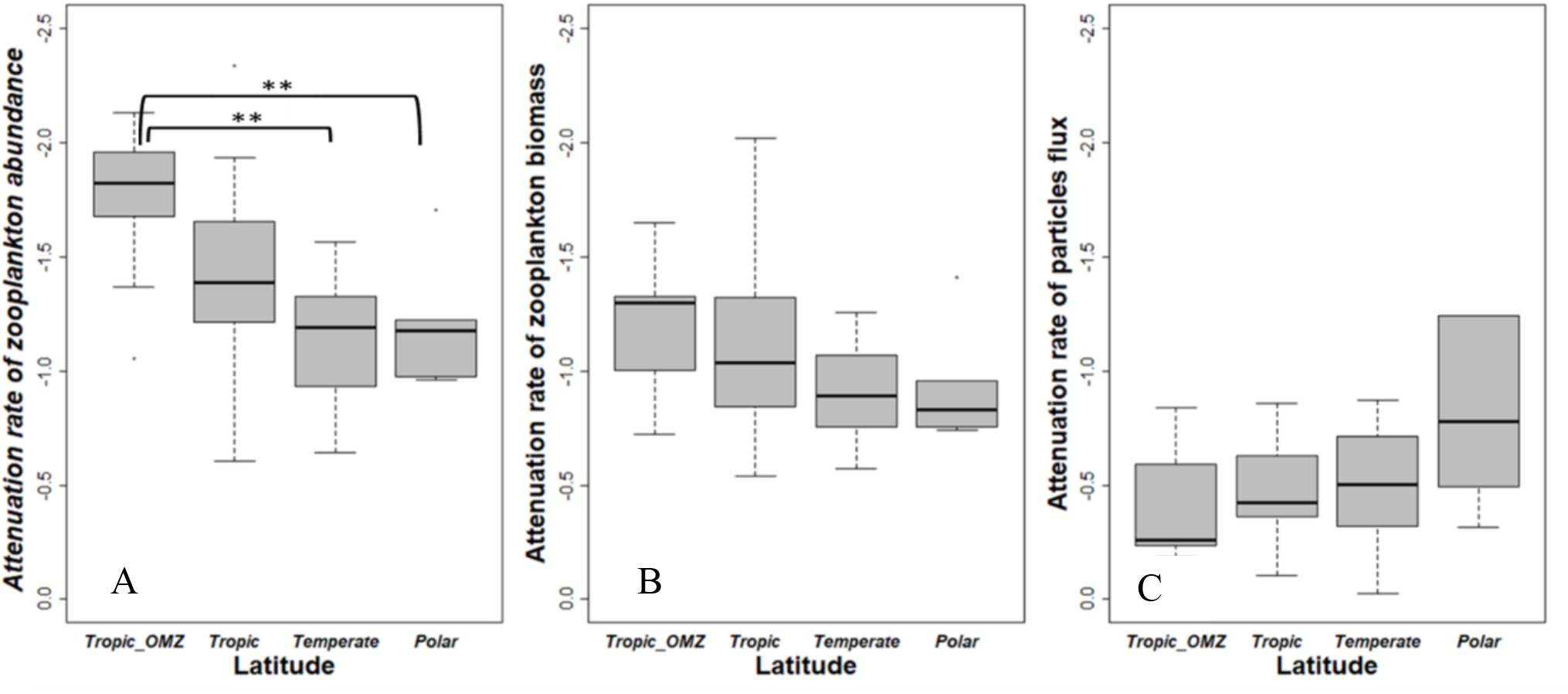
Distributions of attenuation rates at each station per latitudinal bands (Tropical, Tropical OMZ, Temperate, and Polar). A: Attenuation rate of zooplankton abundance (A_zoo). B: Attenuation rate of zooplankton biomass (B_zoo). C: Attenuation rate of particle flux (A_flux). (**) means significant Kruskal-Wallis test with p < 0.01 (Table 5).

**Table 5:** Probability of H0, no difference between the groups (pairwise Kruskal-Wallis test) for total zooplankton and biomass depth attenuation rates and particle flux attenuation rate. * Significant differences.

| Two by two comparisons |  | Abundance | Biomass | Flux |
| --- | --- | --- | --- | --- |
| Tropic-Anoxia | Tropic | 0.057 | 0.641 | 0.77 |
| Tropic-Anoxia | Temperate | 0.0018* | 0.189 | 0.82 |
| Tropic-Anoxia | Polar | 0.01* | 0.238 | 0.09 |
| Tropic | Temperate | 0.34 | 0.639 | 0.99 |
| Tropic | Polar | 0.51 | 0.685 | 0.28 |
| Temperate | Polar | 0.99 | 1 | 0.52 |

### Structuring of the epipelagic community composition

In the epipelagic layer (0 – 200 m depth), the environmental variables explained 32.71% of the total variance in mesozooplankton groups’ abundances. The first RDA axis (RDA1, 57.18 % of constrained variance) opposed the samples from polar waters, and especially those from the Arctic dominated by Calanidae and crustacean larvae (RDA1 > 0), to the tropical samples presenting more even contributions from most of the remaining groups: Protista, Eumalacostraca, Annelida, Amphipoda, Corycaeidae, Chaetognatha, Euphausiacea, Oithonidae, Ostracoda, Oncaeidae, Calanoida (RDA1 < 0). RDA1 was negatively scored by temperature and salinity and positively scored by vertical particle flux, microphytoplankton contribution, suspended particle concentration, dissolved oxygen concentration and chlorophyll *a* concentration. Among the supplementary variables, the attenuations of the particle flux and of the zooplankton biomass were positively correlated with RDA1, while the attenuation of the zooplankton abundance and the Shannon index were negatively correlated to RDA1. RDA2 (13.1% of constrained variance) opposed the samples from the Indian Ocean and North Atlantic Ocean that present higher abundances of Cnidaria, Mollusca, Tunicata and Cladocera (RDA2 < 0) to those samples from the Southern Ocean presenting higher abundances of Annelida, Euphausiacae, Amphipoda and Other Copepoda (RDA2 > 0). RDA2 was positively scored by nitrate concentrations and the relative contribution of nanophytoplankton. It was negatively scored by the concentration of suspended particles and the relative contribution of microphytoplankton. All supplementary variables were negatively correlated with RDA2.

### Structuring of the upper mesopelagic community composition

In the upper mesopelagic layer (200 to 500 m depth), the environmental variables explained 29% of the total variance in mesozooplankton groups’ abundances. Again, the first RDA axis (RDA1, 36.33% of constrained variance) mainly opposed the polar samples dominated by Calanidae copepods, and characterized by higher concentrations of suspended particle, particle flux (both from surface and upper mesopelagic layer), and higher dissolved oxygen concentrations (RDA1 > 0), from the samples characterized by more diverse zooplankton communities (mainly Corycaeidae, small Calanoida, Oncaeidae) and correlated to higher temperature, higher salinity and a higher relative contribution of the nanophytoplankton (RDA1 < 0). Similarly, to what was observed for the epipelagic layer, among the supplementary variables, the attenuations of the particle flux and of zooplankton biomass were positively correlated with RDA1, while the attenuation of zooplankton abundance and the Shannon index were negatively correlated to RDA1. RDA2 (24.72% of constrained variance) opposed the anoxic samples from the Indian Ocean presenting higher abundances of Tunicata, Annelida, Protista, Mollusca, Oithonidae and Cnidaria (RDA2 < 0) to the oxygenated ones displaying higher abundances of Ostracoda, Eumalacostraca, crustacean larvae, other Copepoda, Chaetognatha and Euphausiacea (RDA2 > 0). Samples from the Arctic Ocean were dominated by large copepods from the Calanidae and Metridinidae families. Samples from the Pacific and Atlantic Oceans were dominated by other Copepoda, Eumalacostraca, Ostracoda, Euphausiacea, Chaetognatha and crustacean larvae. Higher attenuation rates of zooplankton biomass and particle flux were found in polar samples whereas higher zooplankton attenuation rates were found in warmer waters, especially at OMZ stations. Again, samples from the tropical upper mesopelagic layers displayed more diverse communities. The distribution of the supplementary variables along RDA2 differed from what was observed for the epipelagic layer, as the supplementary variables, except the attenuation of the zooplankton abundances, were positively correlated to RDA2.

### Structuring of the lower mesopelagic community composition

In the lower mesopelagic layers (500 to 1000 m depth), the environmental variables explained 29.46 % of the total variance in mesozooplankton groups’ abundances. The first RDA axis (RDA1, 35.14 % of constrained variance) opposed the stations from the Arctic and North Atlantic Oceans characterized by higher dissolved oxygen concentrations, higher suspended particle concentrations (RDA1 > 0), from the anoxic and nitrate-rich stations from the Indian Ocean (RDA1 < 0). The former stations were characterized by higher abundances of Calanidae, Calanoida, Metridinidae, crustacean larvae, Ostracoda and Amphipoda, whereas the latter were characterized by higher abundances of Tunicata, Annelida, Eumalacostraca and other Copepoda. RDA2 (22.08 % of constrained variance) separated the colder and less salty stations of the Southern Atlantic Ocean (RDA2 < 0) from the warmer, saltier stations (RDA2 > 0) of the North Atlantic and the Indian Oceans that are characterized by less diverse zooplankton communities. The stations that sampled the oxygenated waters of the North Atlantic and Arctic Oceans were dominated by large copepods of the Metridinidae and Calanidae families, as well as most of the small other Calanoida (i.e. those smaller copepods that could not recognized at a more detailed taxonomic level) in addition to Ostracoda, Amphipoda, and crustacean larvae. Oxygenated stations from the South Atlantic Ocean (SAO) were characterized by higher abundances of Chaetognatha, Corycaeidae, Euphausiacea, Cnidaria, Mollusca, Oithonidae, Oncaeidae and Protista. Samples taken in the OMZ of the Indian and North Pacific Oceans displayed higher abundances of other Copepoda, Tunicata, Annelida, Eumalacostraca and Protista. Higher mesozooplankton biomass and stronger particle flux attenuation rates were found in the normoxic waters while higher zooplankton attenuation rates were observed at the stations that sampled an OMZ. The distribution of the supplementary variables showed similar association with environmental variables as in the upper mesopelagic.

## Discussion

The present global analysis of spatial patterns of the mesozooplankton community and their relationship with the strength of the vertical particle flux is based on zooplankton abundance and biomass estimates and vertical particles flux estimated using state-of-the-art imaging techniques (Zooscan and UVP5). The rigorous quality control (see Supplementary Material) allows to depict with confidence correlations between zooplankton and particle flux, two important components of the biological carbon pump (BCP). Notably the study allows to infer global ecological patterns in the epipelagic and mesopelagic layers, the coupling between these two layers in oxygenated, hypoxic and anoxic situations.

### Important environmental factors for the mesozooplankton community composition in the epipelagic layer

High latitude marine ecosystems are characterised by a combination of lower species diversity and shorter food webs (Laws, 1985; Stempniewicz et al., 2007) sustained by higher concentrations of large microphytoplankton cells (diatoms). On the contrary, low latitude ecosystems are featured by more complex and diverse food webs (Saporiti et al., 2015; Uitz et al., 2006) adapted to lower production rates ensured by smaller cells (i.e. pico- and nanoplankton). How the dynamics of zooplankton community composition and vertical particle flux follow this scheme remains more elusive. Previous field-based studies reported peaks in zooplankton species richness in the tropics (Rombouts et al., 2010; Rutherford et al., 1999; Yasuhara et al., 2012), which is in line with the above-mentioned theory that the tropical food-depleted regions promote more complex food-webs with higher species richness. Our RDAs results result supports the view that, on a global scale, temperature and the production regime of surface ecosystems are the main drivers of zooplankton community structure in the epipelagic layer. Therefore, our observations fall in line with the theory described above: in the epipelagic layers, less abundant and more diverse zooplankton thrive in warm, pico- and nanophytoplankton-dominated waters contrary to the polar waters where zooplankton is much more abundant but less diverse. Polar waters are characterised by a higher contribution of microphytoplankton to total phytoplankton biomass and higher concentrations of particles. Our results also indicate that the two polar communities are not completely alike, as the Arctic is dominated by calanoid copepods while euphausiids and small undetermined copepods dominate in the few stations sampled in the Southern Ocean. Such differentiation has been previously shown by several authors who found that Arctic zooplankton were dominated by *Calanus* spp. (Balazy et al., 2018; Hirche and Mumm, 1992), whereas it was shown that Antarctic zooplankton were dominated by euphausiids, small calanoids, cyclopoids (i.e. *Oithona* spp., *Oncaea* spp.) and salps (Park and Wormuth, 1993; Quetin et al., 1996; Ross et al., 2008).

### Important environmental factors for the mesozooplankton community structure in the mesopelgic layer

The RDA displayed similar general patterns in both the upper and lower mesopelagic layers with the stations in the Indian Ocean (with low zooplankton concentration in low oxygen) and Arctic Ocean (with high zooplankton concentration in oxygenated conditions) being in all layers well separated from the other stations in the remaining ocean. The highest mesopelagic zooplankton concentrations were found at stations associated with high microphytoplankton concentrations in the epipelagic and high particle flux, suggesting a strong impact of surface production regime on the mesozooplankton in the mesopelagic (Hernández-León et al., 2020). The inter-basin differences in zooplankton concentration and community composition was slightly lower in the lower mesopelagic compared to the upper mesopelagic, probably due to the more homogeneous habitat (Fernández de Puelles et al., 2019).

In general, stations associated with anoxic or hypoxic conditions at midwater depth displayed lower zooplankton concentrations in the mesopelagic and different community composition from oxygenated layers. The stations that sampled the tropical OMZ showed higher proportions of gelatinous carnivorous zooplankton (Cnidaria), gelatinous filter feeders (Tunicata), Mollusca and small omnivorous grazers (Cladocera). Anoxic or strongly hypoxic conditions in the mesopelagic may have selected for those taxa adapted to low oxygen (Vaquer-Sunyer and Duarte, 2008). In mesopelagic oxygenated waters, the stations of the Tropical Pacific Ocean displayed a higher diversity stemming from the higher and relatively even abundances of large protists (i.e. Rhizaria and Foraminifera), chaetognaths, crustaceans (Ostracods, Euphausiacea, Eumalacostraca, Amphipoda), and various copepod families (Corycaeidae, Oithonidae, Oncaeidae, small Calanoida). These zooplankton communities were associated with oligotrophic conditions at surface (Fig. 4), lower zooplankton abundances, lower concentrations of suspended matter, weaker particle fluxes and weaker attenuation rates (Fig. 3B) and with stronger attenuation rates of organisms’ abundances (Fig. 3A). Therefore, we evidence oligotrophic regimes where the zooplankton community constitutes a network of diverse taxa that is not as efficient at using the low amount of material fluxing. Our results are consistent with previous studies suggesting that those oligotrophic regimes can be relatively efficient at exporting the slow-sinking fraction of the little carbon that is produced in the surface layers due to a low impact of zooplankton grazing on the sinking particles (Guidi et al., 2009; Henson et al., 2015). This could help explain why we found the lowest particle flux attenuation rates in the tropics.

**Fig. 4:**
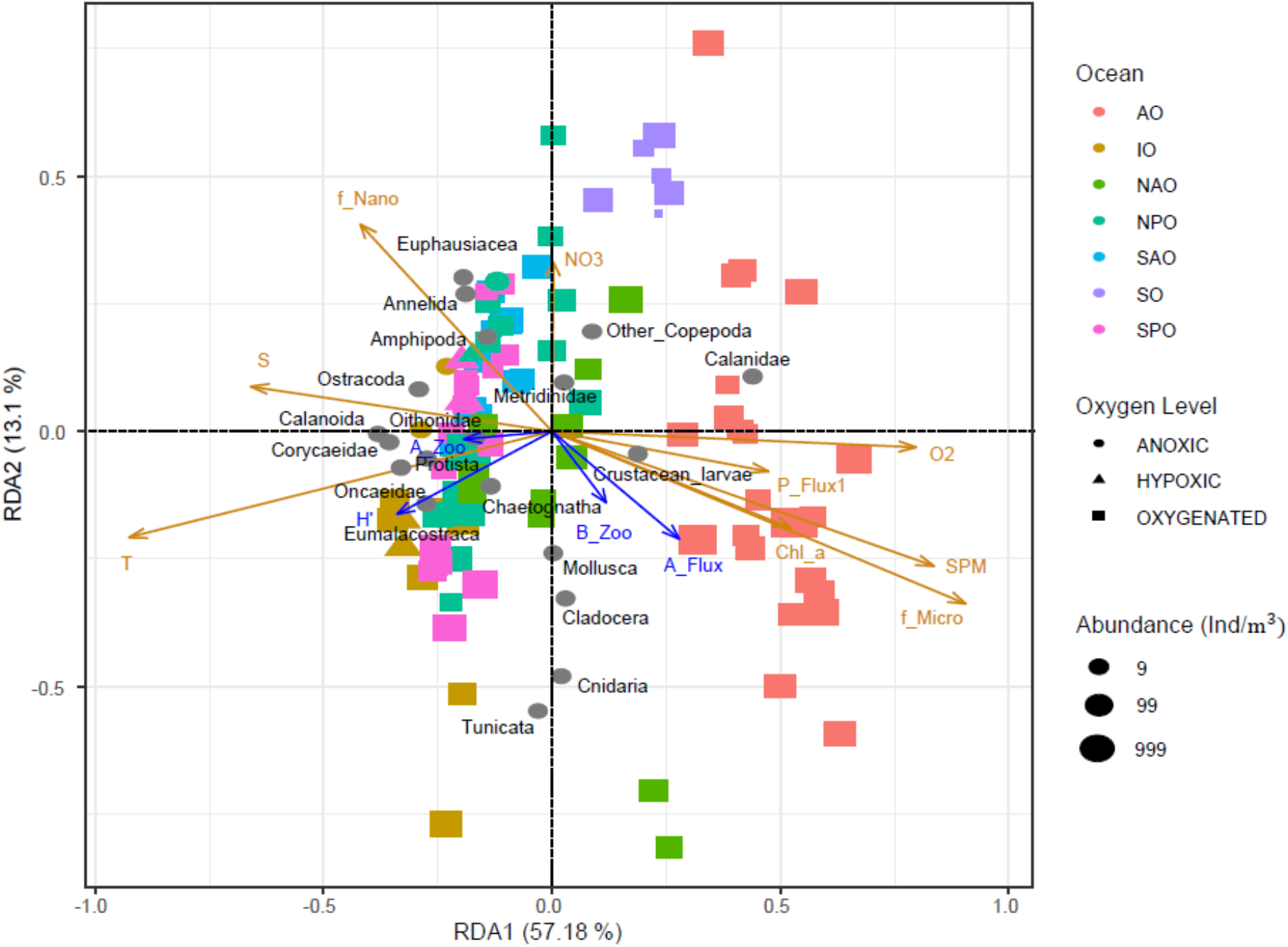
RDA on the epipelagic communities. Each dot corresponds to a sample, i.e., one net at one depth at one station. The orange arrows correspond to the quantitative environmental variables in RDA space: f_pico, f_nano and f_micro correspond to the relative contribution (%) of pico-, nano- and micro-phytoplankton to total phytoplankton biomass, O2 = dissolved oxygen concentration (µmol/kg), Chl_a = Chlorophyll a concentration (mg/m3), SPM = suspended particles matter (m/sr), T = temperature (°C), Sal = salinity, NO3 = nitrate concentration (mol/kg), Z = depth (m)) and P_Flux= particulate flux (*mg*. *m*^-2^. *d*^-1^). Grey dots mark the projection of the zooplankton groups abundance (*ind*. *m*^-3^). Colors correspond to the ocean basin where the samples were taken: AO = Arctic Ocean, IO = Indian Ocean, NAO = North Atlantic Ocean, NPO = North Pacific Ocean, SAO = South Atlantic Ocean, SO = Southern Ocean, SPO = South Pacific Ocean. Shapes illustrate oxygen level, Anoxic: [*O*_2_] <5µmol/kg; Hypoxic: 5µmol/kg< [*O*_2_] <58.5µmol/kg and oxygenated: [*O*_2_]>58.5µmol/k). Supplementary variables estimated for the epipelagic layer are represented with blue arrows: attenuation of particle flux (A_flux), attenuation of zooplankton abundance (A_zoo), attenuation of zooplankton biomass (B_zoo) and the Shannon index (*H’*).

**Fig. 5:**
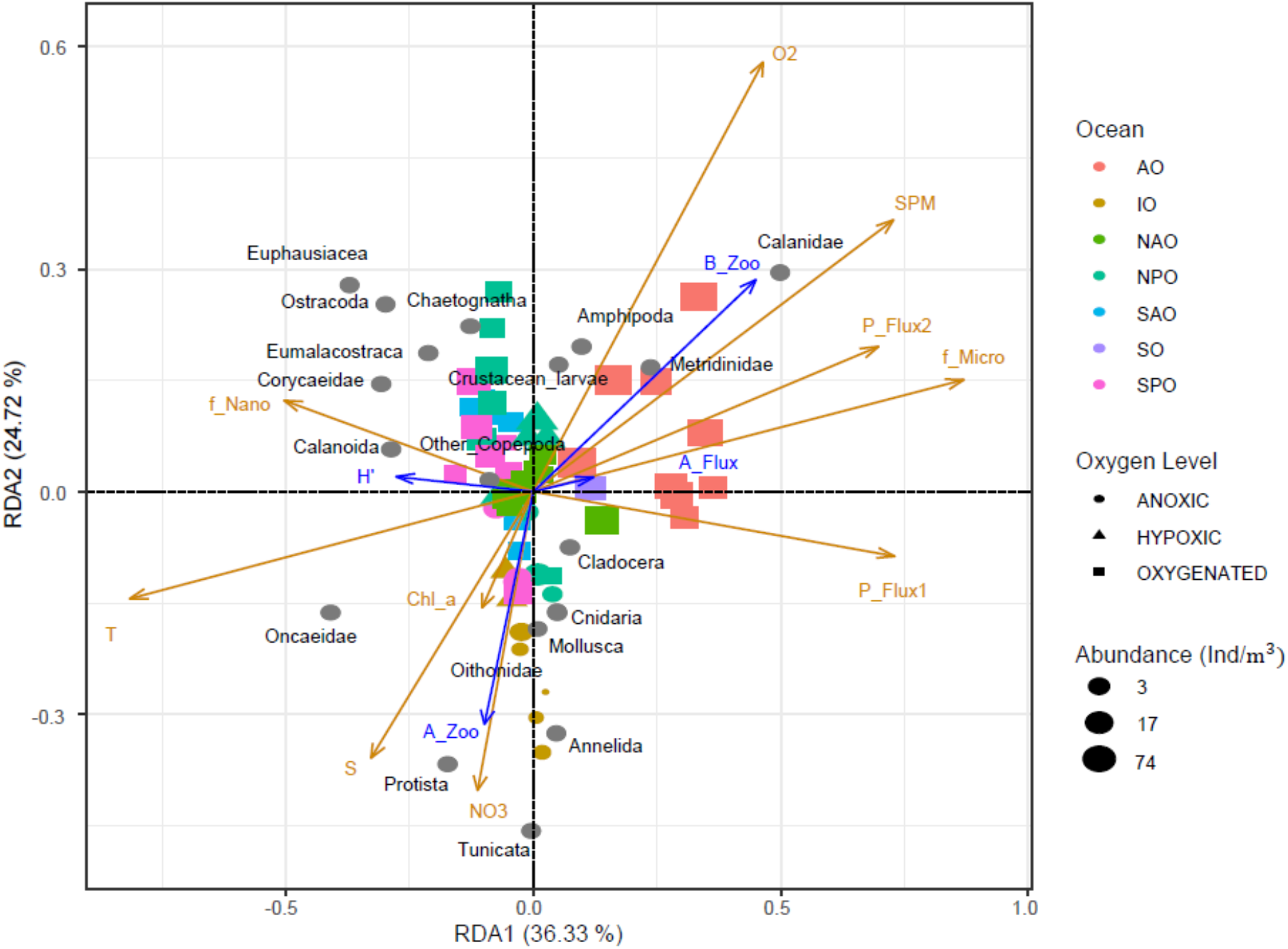
RDA performed on the upper mesopelagic abundances. Each dot corresponds to a sample, i.e., one net at one depth at one station. The orange arrows correspond to the quantitative environmental variables (see legend of Fig. 4). Anoxic: [*O*_2_] <5µmol/kg; Hypoxic: 5µmol/kg< [*O*_2_] <58.5µmol/kg and oxygenated: [*O*_2_]>58.5µmol/kg). Supplementary variables estimated for the upper mesopelagic layer are represented with blue arrows: attenuation of particle flux (A_flux), attenuation of zooplankton abundance (A_zoo), attenuation of zooplankton biomass (B_zoo) and the Shannon index (*H’*).

**Fig. 6:**
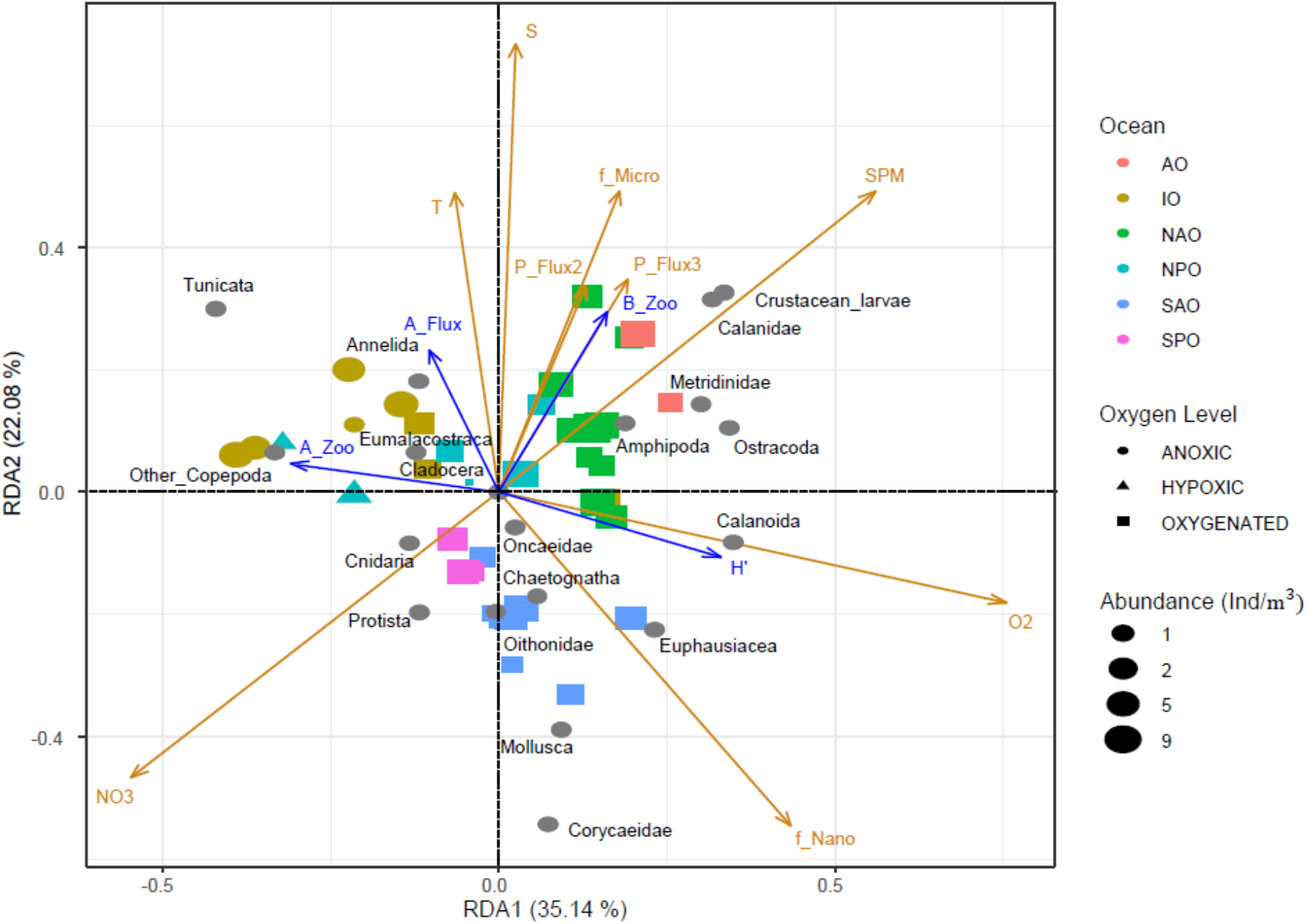
RDA performed on the lower mesopelagic abundances. Each dot corresponds to a sample, i.e., one net at one depth at one station. The orange arrows correspond to the quantitative environmental variables (see legend of Fig. 4). Anoxic: [*O*_2_] <5µmol/kg; Hypoxic: 5µmol/kg< [*O*_2_] <58.5µmol/kg and oxygenated: [*O*_2_]>58.5µmol/kg). Supplementary variables estimated for the lower mesopelagic layer are represented with blue arrows: attenuation of particle flux (A_flux), attenuation of zooplankton abundance (A_zoo), attenuation of zooplankton biomass (B_zoo) and the Shannon index (*H’*).

### Global vertical patterns of zooplankton and particle flux

The concentration and biomass of various zooplankton groups decreased with depth (Fig. S1), confirming the general trend of decreasing zooplankton abundance and biomass from the surface to 1000 m depth resolved (Fig. 2; Bode et al., 2018; Brugnano et al., 2012; Hernández-León et al., 2020; Koppelmann et al., 2005; Kosobokova and Hopcroft, 2010; Yamaguchi et al., 2004). Among these earlier studies, two models for the attenuation of zooplankton have been proposed: an exponential or a power model. Here, we used the power model (equation 1) because it was widely used to model vertical flux attenuation rates (Martin et al., 1987). Based on this power model, we observed rates of decrease in zooplankton abundance or biomass (Fig. 3) that were in the same range as those reported in the western Pacific Ocean (−1.52 to −1.41 for abundance and −1.32 to −1.10 for biomass, (Yamaguchi et al., 2002). Zooplankton abundance decreased more rapidly with depth than biomass as the average size of organisms increased with depth, confirming that the contribution of larger species increases with depth (Homma and Yamaguchi, 2010; Yamaguchi et al., 2002). Considering a larger latitudinal band than earlier studies who found weak regional or global patterns (Hernández-León et al., 2020; Puelles et al., 2019), we showed that the rates of zooplankton attenuation vary with latitude and to a lesser extent with oxygen concentrations (Fig. 3) and also in the opposite direction of the flux attenuation.

Stronger surface particle fluxes and vertical attenuation rates were found at higher latitudes where primary production is mainly ensured by the microphytoplankton and weaker flux and attenuation was found in the low latitude dominated by picophytoplankton (Fig.4). Such latitudinal pattern in particle attenuation rates results from the gradient in the production regime and has been observed quasi-systematically with imaging systems on global scales (Guidi et al., 2015), but also based on sediment traps (Berelson, 2001) or combining sediment traps with satellite-based estimates of primary production (Henson et al., 2012). However, several short-term experiments at selected sites from temperate to tropical latitudes of the Pacific and Atlantic, showed an opposite pattern with stronger flux and attenuation in cold and productive regions compared to inter-tropical oligotrophic (Buesseler et al., 2007; Marsay et al., 2015). The inconsistency of the observations is difficult to explain although it was noted that depth varying remineralisation due to varying temperature may reconcile the contrasted results (Marsay et al., 2015). It is possible that some of the differences arise from the use of different methodologies (instruments, time scale, global representation of the dataset) to address the complexity of the processes to be measured simultaneously.

Such opposite pattern between A_Zoo and B_zoo in one hand and A_Flux in the other hand supports the view that the abundant zooplankton community plays a crucial role in flux attenuation in productive layers, by feeding and fragmenting the sinking material as suggested in previous studies (Dilling and Alldredge, 2000; Lampitt et al., 1990; Sarah L. C. Giering et al., 2014; Stemmann et al., 2004). However, on average and more importantly at high latitudes, zooplankton attenuation rates were stronger than the vertical flux attenuation rates (Fig. 3) indicating that zooplankton vertical zonation may be affected by other processes than the resources provided by the flux of organic matter from the epipelagic layer. In the inter-tropical OMZ regions, zooplankton vertical attenuation rates were maximum when the vertical attenuation of the particle’s flux was minimum. Previous studies have reported lower flux attenuation rates in the OMZ of the Arabian Sea (Haake et al., 1992; Roullier et al., 2014), and Eastern Tropical North Pacific (Cavan et al., 2017; Van Mooy et al., 2002). Lower zooplankton activity together with reduced bacterial activity (Cram et al., 2021) would allow sinking particles to transit through the OMZ core without being severely degraded (Wishner et al., 2008, 1995).

### Sensitivity of plankton and vertical flux to contrasted oxygen conditions

By comparing zooplankton communities from inter-tropical stations with those from oxygenated mesopelagic layers to stations from OMZs, our analysis brought additional evidence that change in zooplankton community composition may affect the efficiency of the BCP. The zooplankton community sampled in the Indian Ocean OMZ displayed a particular composition compared to the other samples, which is in line with the increasing number of studies that document the profound impact of oxygen depletion on pelagic organisms (Ekau et al., 2010; Hauss et al., 2016; Wishner et al., 2018). Our results underlined how the OMZ can strongly reduce the abundance of several zooplankton groups (i.e. Calanoida, Euphausiacea, Amphipoda, Ostracoda) that are outcompeted by more hypoxia-tolerant ones. We found that tunicates (mostly Appendicularia), large protists (Collodaria and Foraminifera), polychaetes, Oncaeidae, Oithinidae and to a lesser extent Cnidaria (jellyfishes) may tolerate OMZ conditions, since their abundance at OMZ and at non-OMZ stations did not present significant differences. All of these zooplankton groups have been reported as being able to thrive or endure at low oxygen concentrations in various OMZs (Ekau et al., 2010; Hauss et al., 2016; Keister and Tuttle, 2013; Kiko and Hauss, 2019; Parris et al., 2014; Tutasi and Escribano, 2020; Werner and Buchholz, 2013; Wishner et al., 2020). Those “hypoxiphilic” or hypoxia-tolerant taxa display special adaptations that enable them to remain at extremely low oxygen concentrations where other zooplankton groups cannot, because they fail to meet their metabolic oxygen demand (Childress and Seibel, 1998; Seibel, 2011). Jellyfish and large protists benefit from their passive feeding tactics which are less oxygen-demanding than active cruising and filter feeding (Kiørboe, 2011). Indeed, hypoxia-tolerant taxa can also display behavioral adaptations that cut down the metabolic costs associated with active feeding or the active search for mates in the water column. Copepod species belonging to the Oncaeidae and Oithonidae families are known for performing ambush feeding tactics or to attach themselves on large detritus (Brun et al., 2017), which allows them to capture small preys or to feed on falling detritus at very low metabolic costs. Therefore, these less motile copepods frequently outcompete most of their calanoid congeners in OMZs and often co-occur with Appendicularia, as those discard particle-rich aggregates on which the copepods feed (Alldredge, 1972; Brun et al., 2017; Kiørboe, 2011). Other possible adaptations consist in the use of lipid storages and metabolic suppression(Wishner et al., 2020). Some organisms conduct a diapause in the OMZ for an extended period, allowing them to avoid foraging predators and to complete their life cycle (Arashkevich et al., 1996). Diapause reduces energetic costs and thus allows the organisms to survive in resource-depleted conditions. Metabolic suppression goes hand in hand with the diapause, but can also occur on shorter time scales, e.g. when organisms stay at OMZ depth during diel vertical migrations. Diapause and metabolic suppression are associated with reduced respiratory and locomotor activity. Feeding on and disruption of particles could therefore be reduced at OMZ depth not only due to the exclusion of many hypoxia intolerant zooplankton organisms, but also due to the reduced activity of those zooplankton organisms that can cope with OMZ conditions. The zooplankton groups we found to characterize the community inhabiting the hypoxic and anoxic layers are those that commonly outcompete others in OMZs worldwide. They could benefit from the likely future expansion of OMZ (Oschlies et al., 2018) and thus become increasingly abundant with future climate change. This would probably result in enhanced vertical particle flux in those regions. However, this requires further research as some of these organisms could already be living near their physiological hypoxia tolerance limits (Wishner et al., 2018).

## Conclusions

Our study combined consistent and large-scale observations of zooplankton abundance, biomass and community composition with estimates of vertical particle fluxes obtained through imaging system pipelines. We showed that consistent observations can be obtained at a global level using in situ camera systems and precise net sampling in various ecosystems representative of different ocean conditions. In future surveys, consistently combining these techniques with acoustic and other bio-optical sensors will allow the measurement of the vertical and horizontal distribution of organisms with greater precision. We showed that the key environmental variables driving epipelagic mesozooplankton community structure at the global scale are temperature, phytoplankton biomass and phytoplankton key groups. In the mesopelagic layer, surface phytoplankton size classes, particle concentration, temperature and oxygen availability were identified as the main drivers of mesozooplankton community structure. Our work furthermore suggests that low attenuation of zooplankton abundance and biomass go in hand with high particle flux attenuation and vice versa. Such information is crucial for the parameterization of the next generation of marine ecosystem models that describe complex zooplankton-related processes based on coarse but increasingly numerous functional types. Such models suggest that surface phytoplankton biomass and size classes, the flux of particles and the oxygen content in the mesopelagic layer will all be affected further by global climate change. The fact that zooplankton communities are sensitive to those factors suggests that future climatic changes may profoundly alter zooplankton communities worldwide, at the surface but also in the deeper mesopelagic layer.

## ACKNOWLEDGMENTS

Tara Oceans (which includes both the Tara Oceans and Tara Oceans Polar Circle expeditions) would not exist without the leadership of the Tara Ocean Foundation and the continuous support of 23 institutes (https://oceans.taraexpeditions.org/). This study is part of the “Ocean Plankton, Climate and Development” project funded by the French Facility for Global Environment (FFEM). Y.D.S. and M.C.B. received financial support from FFEM to execute the project. R.K acknowledges support support via a “Make Our Planet Great Again” grant of the French National Research Agency within the “Programme d’Investissements d’Avenir”; reference “ANR-19-MPGA-0012”. F.B. received support from ETH Zürich and the European Union’s Horizon 2020 research and innovation programme under grant agreement n°SEP-210591007. We further thank the commitment of the following sponsors: CNRS (in particular Groupement de Recherche GDR3280 and the Research Federation for the Study of Global Ocean Systems Ecology and Evolution FR2022/Tara Oceans-GOSEE), the European Molecular Biology Laboratory (EMBL), the French Ministry of Research, and the French Government ‘‘Investissements d’Avenir’’ programs OCEANOMICS (ANR-11-BTBR-0008), the EMBRC-France (ANR-10-INBS-02). Funding for the collection and processing of the Tara Oceans data set was provided by NASA Ocean Biology and Biogeochemistry Program under grants NNX11AQ14G, NNX09AU43G, NNX13AE58G, and NNX15AC08G (to the University of Maine); the Canada Excellence research chair on remote sensing of Canada’s new Arctic frontier; and the Canada Foundation for Innovation. We also thank Agnès b. and Etienne Bourgois, the Prince Albert II de Monaco Foundation, the Veolia Foundation, Region Bretagne, Lorient Agglomeration, Serge Ferrari, Worldcourier, and KAUST for support and commitment. The global sampling effort was enabled by countless scientists and crew who sampled aboard the Tara from 2009–2013, and we thank MERCATOR-CORIOLIS and ACRI-ST for providing daily satellite data during the expeditions. We are also grateful to the countries who graciously granted sampling permission.

## Supplementary material

### Availability of the dataset

**Data sets for the environment are available at Pangea:**

(https://doi.org/10.1594/PANGAEA.840721)

**Tables for the RDA are available**

For zooplankton: Metadata_Zooplankton_Article_RDA.xls

**Zooplankton Pangea for the tables/ecotaxa for the vignettes**

https://ecotaxa.obs-vlfr.fr/prj/714?taxo=56693&taxochild=N&ipp=100&zoom=100&sortby=&magenabled=0&popupenabled=0&statusfilter=&samples=&instrum=&sortorder=asc&dispfield=&projid=714&pageoffset=0

### Quality of the dataset

The analysis reported here is the first to apply consistent imaging techniques in order to investigate particle flux, zooplankton community composition and biomass across the global ocean. To do so, our study relies on the combined use of the Underwater Video Profiler v5 (UVP5), a Multinet and the Zooscan imaging system. These results are the first combined global zooplankton and flux estimates and as such provide a baseline for future studies. Their quality needs to be discussed with regards to 1) flux assessment, 2) mesh selectivity, mass conversion and possible impact of dead zooplankton in the images, the capacity of the sampling design to capture or not 3) day night variability and 4) mesopelagic peaks of zooplankton.

Vertical particle mass fluxes (mg Dry Weight m^-2^d^-1^) were calculated from the particle size spectra (150 µm −1.5 mm) detected by the UVP5 following Guidi et al., (2008). Briefly, the procedure optimized two parameters in the flux equation by comparing UVP5 derived flux and a global sediment trap flux data set (Guidi et al., 2008). Depending on the ocean basin, regional algorithms with different sets of the two parameters have been proposed (Forest et al., 2013; Kiko et al., 2017) but the shape of the allometric relationships between particle size and flux is kept as a growing monotonic power relationship. Many studies have shown that this assumption is verified when comparing extremes size of particles but they all have also shown a high variability (Guidi et al., 2008; Iversen and Lampitt, 2020) which cannot be depicted without more information on the type of particles. In addition, we used the same parametrization for each depth layer although a few observational and modelling studies (Berelson, 2002; Gehlen et al., 2006) have proposed that the sinking speed may vary with depth. Finally, the fluxes are calculated considering all particles detected in situ by the UVP5. Because particles are so abundant in the size range presently considered for flux estimation (150µm to 1.5mm), the impact of zooplankton in the present flux estimates is low (Guidi et al., 2015; Kiko et al., 2020). We did not perform further sensitivity analyses as the aim of this study was not to quantify absolute carbon flux but to assess correlations with zooplankton and other environmental variables. In the future, better recognition of particles on images will possibly allow to calculate different particle speed for different ypes of aggregates as was proposed recently (Trudnowska et al., 2021).

Zooplankton abundance estimates are sensitive to the mesh of the plankton net tow used (Gallienne et al., 2001). A mesh size of 300µm was selected for the Multinet as a good compromise to capture both mesozooplankton and macro-zooplankton. However, the small copepod families (Oithona, Oncaeidae, Clausocalanidae, Paracalanidae) that dominate community composition in tropical/subtropical open oceans are not collected in an optimal fashion with the present mesh size, contrary to the larger Calanidae notably in the temperate and polar regions. Therefore, our abundance estimates likely underestimate the contribution of the smallest mesozooplankton groups. Nonetheless, even a net equipped with a 150 µm mesh can still underestimate the abundance of cyclopoids and early copepod stages (Paffenhofer and Mazzocchi, 2003).

Nets can also destroy fragile organisms during the sampling process (Remsen et al., 2004). For example, the very low abundance and biomass of Rhizarians reported here contrasting with recent findings based on *in situ* cameras (Biard et al., 2016). This is likely due to such sampling biases. As for the estimation of particle fluxes from particle size spectra, we assessed the carbon biomass of each individual organism’s size using published allometric relationships (Lehette and Hernández-León, 2009) in which large and rare organisms contributes importantly to the total biomass bringing strong variability in the results. To account for the excessive variability in biomass estimates, we analysed our results using both abundance and biomass.

Dead zooplankton, including carcasses, are increasingly recognized as being quite abundant in the mesopelagic water (Böttger-Schnack, 1996; Roe, 1988; Terazaki and Wada, 1988; Wheeler Jr, 1967). It is impossible to assess whether the damages of the organisms observed on the digitized organisms were inflicted by the net tow. This is even harder to assess in the deeper nets which spent more time in the water column. Therefore, we cannot tell whether the organisms were damaged during the sampling process or dead beforehand. We attempted to separate the carcasses from the complete bodies of copepods based on grey intensity criteria or by looking at the physical integrity of the organisms, but this was not successful because of independent and continuous changes for both criteria. Nonetheless, among the 110’000 vignettes of copepods, 21’117 (20%) were initially classified as possible carcasses. Their proportion was equal to 42.90% in the surface layer, 26.71% in the upper mesopelagic and 30.39% in the deepest layers. The median and interquartile of abundances varied from the surface (median = 1.21 *Ind*/*m*^3^, IQR = 2.53) to the upper mesopelagic (median = 0.75, IQR = 0.89) and increased towards deeper layers (median = 0.85, IQR = 0.99). Subsequent data analyses showed that aggregating them or not with the non-damaged copepods had no effect on the analysis’ results because their proportion was homogeneous across sampling sites. Therefore, all copepods were ultimately pooled together for all our numerical analyses.

Zooplankton and micronekton perform diel vertical migration (DVM) in all oceans (Banse, 1964). Depending on the organism’s taxonomic group and size, ranges of diel migration show tremendous variability (Ohman and Romagnan, 2016; Tarling et al., 2004). In our study, significant differences in abundances between day and night samples was found for a few groups known to be diel migrators (Euphausiids, Metridinidae, Corycaeidae, Cnidaria for abundance data; and Eumalacostraca and Ostracoda for biomass data). However, at the community level (i.e., total mesozooplankton), no significant differences could be found between the day and night samples at surface. However, total zooplankton biomass was higher in night samples compared to day samples, because of the more abundant large-bodied migrators (Eumalacostraca, Euphausiacea, Table S2). For abundance, this is a rather unexpected result as a large number of zooplankton taxa perform DVM (Kiørboe et al., 2018; Ohman and Romagnan, 2016). Yet, this absence of significant changes had already been observed in previous studies using similar types of plankton nets (Wishner et al., 2018). DVM could not be detected from the ANOSIM analysis of the surface layer for different reasons. First, the weak diel differences in abundances could easily be smeared by plankton patchiness when the replicates are too low. Second, it is possible that no differences were observed because the large organisms carrying out DVM are too few in number compared to the smaller ones in the present data. Day and night differences between the slopes of the total biomass were found, probably due to the few diel migrators species. Hence, it is well known that large zooplankton (euphausiids, sergestid shrimps, amphipods, fish such as myctophids(Morrison et al., 1999)) sampled with larger Multinet nets (opening >1m^2^ and mesh size >500µm), or detected through acoustics techniques, showed stronger DVM than smaller organisms collected through smaller Multinet nets such as ours (opening of 0.25 m^2^ and mesh size of 300µm). Furthermore, copepods contributed to nearly 80% of total zooplankton abundance in our samples, and copepod DVM may be confined most typically to populations living within the epipelagic zone (Madhupratap et al., 2004; Morrison et al., 1999; Smith et al., 1998).

Many studies have reported midwater peaks of zooplankton and nekton in 100-200 m depth layers centered around 40 to 600 m depth (Andersen et al., 1998; Irigoien et al., 2014; Koppelmann et al., 2004) indicating that the use of a simple power law to describe plankton biomass may be an oversimplification of the true vertical zonation. The general lack of such peaks may be explained by the variability in methods used between studies. Midwater peaks are often reported with instruments allowing a good vertical resolution (> 10 observations in the upper kilometer) and targeting the large migrating organisms with nets having large mesh size or acoustic devices with frequencies adapted to nektonic organisms. Many studies using Multinets equipped with rather fine mesh size (<500µm) do not capture such peaks in abundance/ biomass (Homma and Yamaguchi, 2010; Yamaguchi et al., 2002). In our study, the mesopelagic layer was sampled down to 900 m depth, and only with 2 or 3 nets with a mesh size of 300µm and a small net opening (0.25 m^2^). Therefore, midwater peaks could have been smoothed in the few layers. Hence, the power law simplification may be a proper proxy for the smaller fraction of the mesozooplankton (mainly copepods smaller than 2 mm).

### Community vertical distribution

On a global scale, zooplankton average abundance and biomass decreased drasctically with depth, with quasi similar patterns for the eight main zooplankton groups (Fig. S1). The strongest variation in abundance was observed for copepods, followed by chaetognaths, small omnivorous zooplankton and crustacean decapods (Fig. S1). The strongest variation in biomass with depth were observed for copepods followed by decapods, chaetognaths and small omnivorous zooplankton (Fig. S2).

**Fig. S1:**
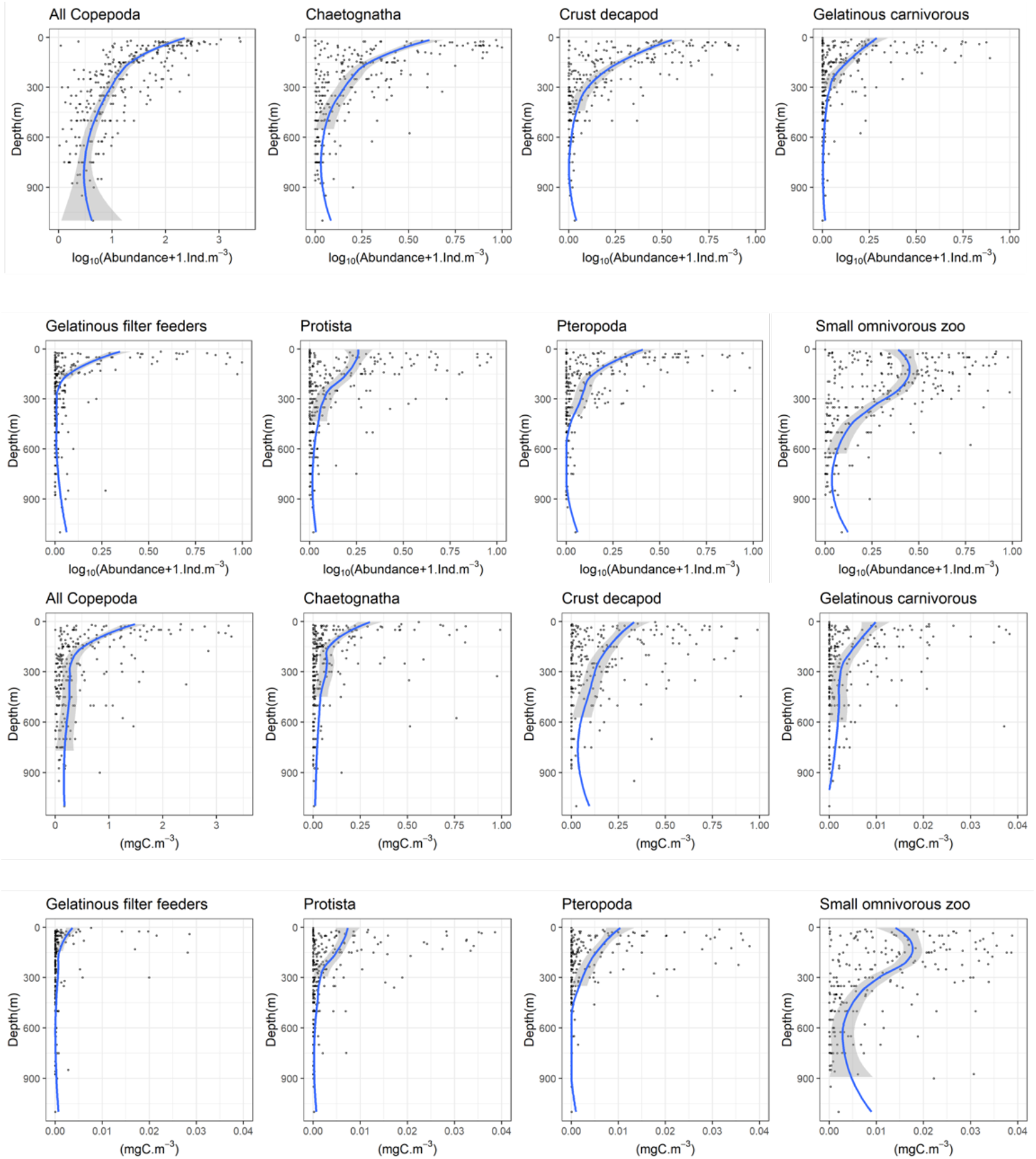
Vertical profiles of abundance (two first lines) and biomass (two last lines) in the global ocean for the 8 functional mesozooplankton groups: all copepoda, chaetognatha, crustacean decapoda (crust decapod), gelatinous carnivorous, gelatinous filter feeders, protista, pteropoda, small omnivorous zooplankton. Not all abundance and biomass scales are similar to better observe the gradients. In all cases, solid lines correspond to a smooth trends and gray ribbons to the 95% confidence intervals. These trends are drawn for illustrative purposes and were not used in down-stream analyses.

### Community composition in the upper first kilometer of the ocean

When analyzing all vertical layers together, the redundant discriminant analysis (RDA) indicated that environmental variables explained 27.01% of the variance in mesozooplankton groups abundance (Fig. 4). The first RDA axis (RDA1; 47.08% of the variance constrained by the RDA) opposed the samples from polar waters (Arctic Ocean, Southern Ocean; RDA1 > 0) to the samples taken at lower latitudes (Indian Ocean, Northern and Southern Atlantic Ocean, Northern and Southern Pacific Ocean; RDA1 < 0), especially for epipelagic and upper mesopelagic samples. RDA1 was negatively scored by variables depicting a gradient of tropical to extratropical conditions: temperature, salinity, and the relative contribution of picophytoplankton and nanophytoplankton to phytoplankton community biomass. Meanwhile, RDA1 was positively scored by dissolved oxygen concentration, the intensity of the vertical particle flux and its attenuation rate, the concentration of suspended particles and the contribution of microphytoplankton to total phytoplankton biomass. The second RDA axis (22.22 % of the constrained variance) opposed the samples from the surface (RDA2 < 0) to the samples from mesopelagic layers (RDA2 > 0). This axis was positively scored by depth, salinity, and nitrate concentrations, and negatively scored by pico-, nano- and microphytoplankton concentrations, oxygen concentration, particles and chlorophyll *a* concentrations. Samples from the epipelagic layer of higher latitudes showed higher plankton abundances and were dominated by Calanidae and crustacean larvae (copepod nauplii). Samples from the epipelagic layers of lower latitudes Tropical Ocean were more diverse as they displayed more even contributions of Protista, Eumalacostraca, Annelida, Amphipoda, Tunicata, Corycaeidae, Chaetognatha, Euphausiacea, Oithonidae, Cnidaria, Mollusca and Cladocera. In the mesopelagic layer, communities were less spread in the RDA space than for the surface layer. Mesopelagic communities were dominated by Other Copepoda and Metridinidae at higher latitudes and by Calanoida, Ostracoda and Oncaeidae at lower latitudes. Polar and temperate waters displayed lower mesozooplankton diversity but higher mesozooplankton abundances/biomass, more intense vertical particle fluxes, and stronger particle attenuation rates. On the opposite, the warmer and more oligotrophic tropical waters displayed lower diversity, lower mesozooplankton biomass/ abundance, but higher attenuation rates of mesozooplankton organisms. To better disentangle the relationships between mesozooplankton composition and the environmental conditions within the different vertical layers of the ocean, separate RDAs were then performed for each depth layer.

**Figure S2:**
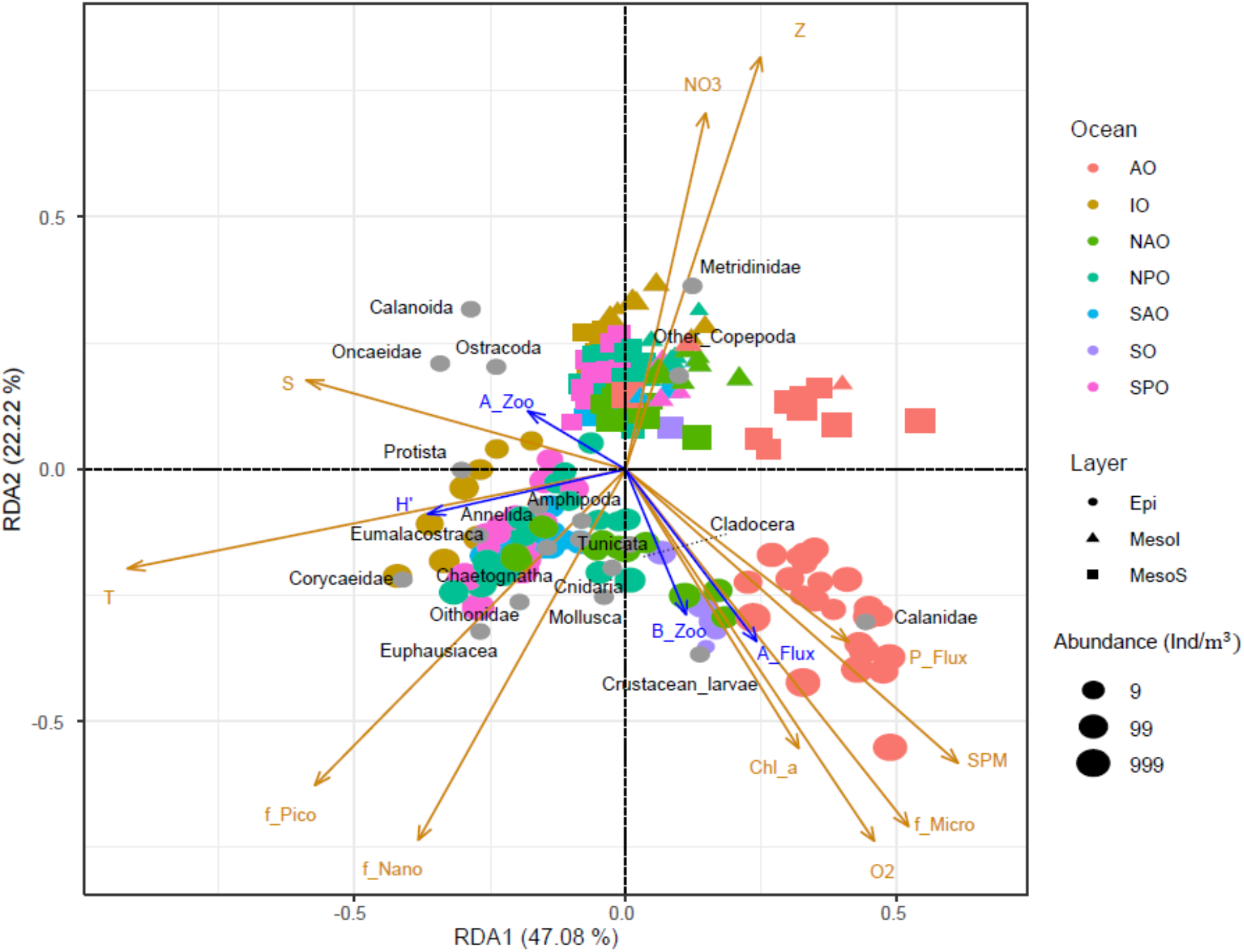
RDA performed with all layers. Each colored dot corresponds to a sample, i.e., one net at one depth at one station. The orange arrows mark the projection of the environmental variables in RDA space: f_pico, f_nano and f_micro correspond to the relative contribution (%) of pico-, nano- and micro-phytoplankton to total phytoplankton biomass, median value of (O2 = dissolved oxygen concentration (µmol/kg), Chl_a = Chlorophyll a concentration (mg/m3), SPM = suspended particles matter (m/sr), T = temperature (°C), Sal = salinity, NO3 = nitrate concentration (mol/L), Z = depth (m)) and P_Flux= particulate flux (*mg*. *m*^-2^. *d*^-1^)). Grey dots mark the projection of the zooplankton groups abundance (*ind*. *m*^-3^). Colors correspond to the ocean basin where the samples were taken: AO = Arctic Ocean, IO = Indian Ocean, NAO = North Atlantic Ocean, NPO = North Pacific Ocean, SAO = South Atlantic Ocean, SO = Southern Ocean, SPO = South Pacific Ocean. Shapes illustrate the vertical layers where the samples were taken: Epi = epipelagic, MesoS = upper mesopelagic and MesoI = lower mesopelagic.

**Table S1:** List of nets that were removed from the RDA analysis. AO=Arctic Ocean, IO=Indian Ocean, NAO=North Atlantic Ocean, NPO=North Pacific Ocean, SAO=South Atlantic Ocean, SO=Southern Ocean or Austral Ocean, SPO=South Pacific Ocean.

| Ocean Bassin | Nets |
| --- | --- |
| IO | 36_4,38_4, 39_2, 42_3 |
| AO | 163_2, 163_3, 206_2 |
| NAO | 142_4,142_4n,143_3,143_3n,147_3,147_4,150_1,150_4,151_2,151_4,152_2,152_3,155_2,158_2 |
| NPO | 131_2, 132_2,133_3,135_2,137_3,137_3n |
| SAO | 131_2, 132_2,133_3,135_2,137_3,137_3n |
| SO | 84_1, 85_1, 85_2 |
| SPO | 98_2, 98_3, 102_1, 106_2,<br>106_4,109_1,110_1,110_1n,111_1,111_1n,122_1,122_1n,125_1,125_3,127_2,127_2n |

**Table S2:** Wilcoxon test for day and night for each taxon at level of 5% (bold values mean significant test)

| Species | Wilcoxon p-value (abundance) |  | Wilcoxon p-value (biomass) |  |
| --- | --- | --- | --- | --- |
|  | Epipelagic | Upper Mesopelagic | Epipelagic | Upper Mesopelagic |
| Protista | 0.50 | 0.13 | 0.8 | 0.06 |
| Annelida | 0.66 | 0.82 | 0.62 | 0.57 |
| Chaetognatha | 1 | 0.20 | 0.58 | 0.32 |
| Tunicata | 0.26 | 0.078 | 0.06 | 0.19 |
| Cnidaria | 0.90 | **0.037** | 0.58 | **0.009** |
| Mollusca | 0.95 | 0.123 | 0.26 | 0.96 |
| Eumalacostraca | 0.39 | 0.62 | **0.049** | 0.49 |
| Euphausiacea | 0.21 | **0.0098** | **0.0001** | 0.63 |
| Amphipoda | 0.07 | 0.91 | 0.39 | 0.83 |
| Ostracoda | 0.32 | 0.96 | **0.049** | 1 |
| Crustacean_larvae | **0.01** | 0.81 | 0.23 | 0.81 |
| Other_Copepoda | 0.76 | 0.46 | 0.95 | 0.41 |
| Calanoida | 0.10 | 0.24 | 0.11 | 0.32 |
| Oithonidae | 0.19 | 1 | 0.24 | 0.84 |
| Corycaeidae | **0.01** | 1 | **0.016** | 0.76 |
| Oncaeidae | 0.66 | 0.41 | 0.8 | 0.51 |
| Metridinidae | **0.0001** | **0.018** | **0.0001** | **0.006** |
| Calanidae | 0.29 | 0.12 | 0.07 | 1 |

## Notes

### Competing Interest Statement

The authors have declared no competing interest.

